## Supplementary Information for "Multistable and dynamic CRISPRi-based synthetic circuits"

| Name | Addgene ID | Resistance | Ori | Relevant features | Related figure(s) |
|---|---|---|---|---|---|
| pC-0 | 124421† | Kanamycin | ColA | Empty multiple cloning site | Used for non-fluorescent control bacteria |
| pC-0_v2 | 124422† | Ampicillin | ColA | Empty multiple cloning site | Used for non-fluorescent control bacteria |
| pJ1996_v2 | 140664† | Spectinomycin | CloDF13 | dCas9 & Csy4 | Fig. 1, Fig. 4, Fig. 5, Suppl. Fig. 1, Suppl. Fig. 2, Suppl. Fig. 3, Suppl. Fig. 4, Suppl. Fig. 6 |
| pJ2018 | 140665 | Spectinomycin | CloDF13 | dCas9, LuxR & Csy4 | Fig. 2 |
| pJ2077.2 | 140666 | Ampicillin | pSC101 | Controller plasmid for TS circuit and cL and cR controls | Fig. 2 |
| pJ2076.2_TS | 140667 | Kanamycin | ColA | Toggle switch (TS) | Fig. 2 |
| pJ2076.2_cL | 140668 | Kanamycin | ColA | cL control for toggle switch experiment | Fig. 2 |
| pJ2076.2_cR | 140669 | Kanamycin | ColA | cR control for toggle switch experiment | Fig. 2 |
| pJ2042.2 | 140670 | Kanamycin | ColA | 3-color stripe | Fig. 4a, b, c & d, Suppl. Fig. 1 |
| pJ2042.2_Bs | 140671 | Kanamycin | ColA | 1-color stripe ("BLUE") | Fig. 4e, f, g & h, Suppl. Fig. 3 |
| pJ2048.2_Gs | 140672 | Ampicillin | ColA | 1-color stripe ("GREEN") | Fig. 4e & f, Suppl. Fig. 3 |
| pJ2048_2xNOT | 140673 | Ampicillin | ColA | Double-inverter | Fig. 4g & h |
| pJ2072.2_CRISPRlator | 140674 | Kanamycin | ColA | CRISPlator | Fig. 5 |
| pJ2072.2_c1 | 140675 | Kanamycin | ColA | CRISPlator control ("open ring") | Suppl. Fig. 6 |
| pJ2044 | 140676 | Kanamycin | ColA | NOT gate with sg-1 | Fig. 1 |
| pJ2043 | 140677 | Kanamycin | ColA | NOT gate with sg-2 | Fig. 1 |
| pJ2039 | 140678 | Kanamycin | ColA | NOT gate with sg-3 | Fig. 1 |
| pJ2040 | 140679 | Kanamycin | ColA | NOT gate with sg-4 | Fig. 1 |
| pJ2044_N2only | 140680 | Kanamycin | ColA | Control ("C") lacking sg-1 | Fig. 1 |
| pJ2043_N2only | 140681 | Kanamycin | ColA | Control ("C") lacking sg-2 | Fig. 1 |
| pJ2039_N2only | 140682 | Kanamycin | ColA | Control ("C") lacking sg-3 | Fig. 1 |
| pJ2040_N2only | 140683 | Kanamycin | ColA | Control ("C") lacking sg-4 | Fig. 1 |
| pJ2044_t4 | 140684 | Kanamycin | ColA | NOT gate with sg-1t4 | Fig. 1 |
| pJ2040_t4 | 140685 | Kanamycin | ColA | NOT gate with sg-4t4 | Fig. 1 |
| pJ2042.2_GFPonly | 140686 | Kanamycin | ColA | 1-color stripe | Suppl. Fig. 2a |
| pJ2048.2 | 140687 | Kanamycin | ColA | 3-color stripe (different sgRNAs) | Suppl. Fig. 2b |
| pJ2042.2_invRep | 140688 | Kanamycin | ColA | 3-color stripe (inverted reporters) | Suppl. Fig. 2c |
| pJ2042.2_Brocc | 140689 | Kanamycin | ColA | Broccoli stripe | Suppl. Fig. 4 |

**Supplementary Table 1.** Plasmids used in this study. †Plasmids from Santos-Moreno & Schaerli 2019, ACS Synth. Biol. 8, 1691-1697. The rest of the plasmids were constructed in this study.

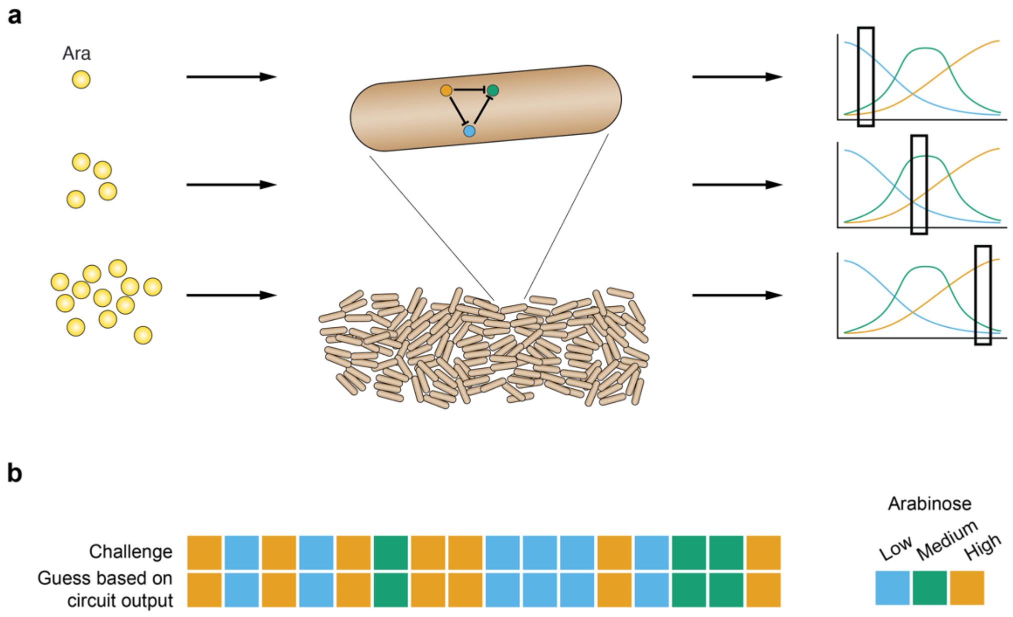


**Supplementary Fig. 1.** Bacteria carrying the CRISPRi stripe network (Fig. 4) work as a biosensor for Ara concentration determination. (**a**) Engineered bacteria bearing the IFFL circuit exhibit distinct levels of the three fluorescent reporters depending on the Ara concentration they are subjected to. (**b**) 16 blinded solutions with low, medium or high Ara were used to challenge the engineered bacteria, and the identity of the challenge was identified with 100% accuracy based on the fluorescence readout. Data from three biological replicates.

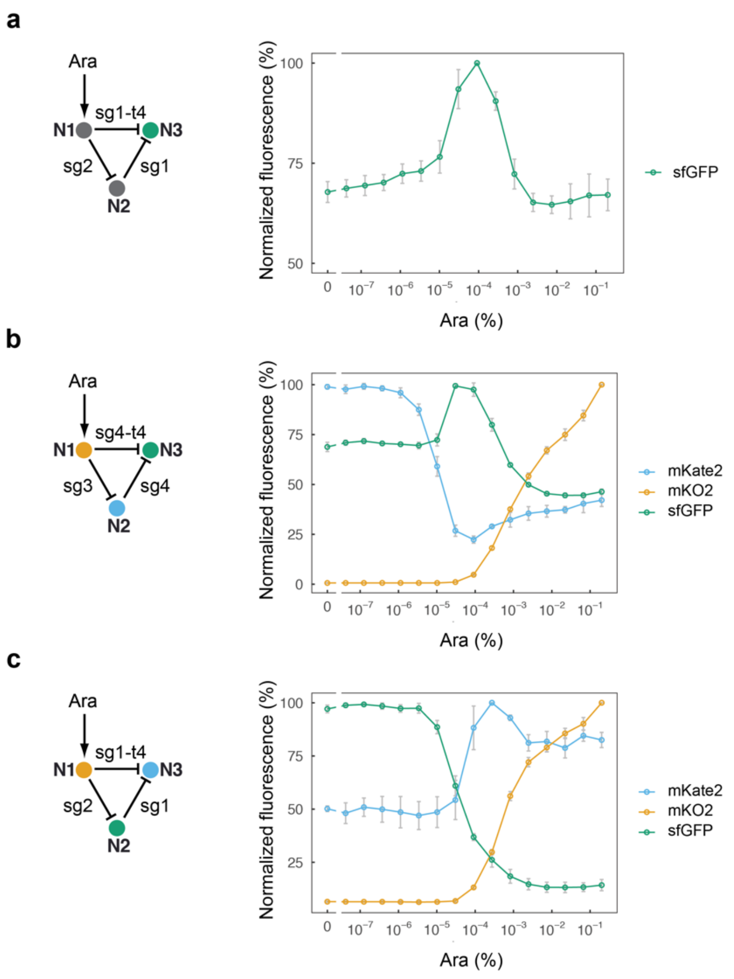


**Supplementary Fig. 2.** Robustness of the CRISPRi IFFL to variations in its design. (**a**) Design and behavior of a circuit with only one reporter (sfGFP-MarAn20 in N3) instead of three. (**b**) Design and behavior of a network relying on a different set of regulators. (**c**) Design and behavior of a circuit in which reporters for N2 and N3 are inverted compared to the original design. All networks still display a stripe and thus showcase the robustness of the design. Mean and s.d. of three biological replicates.

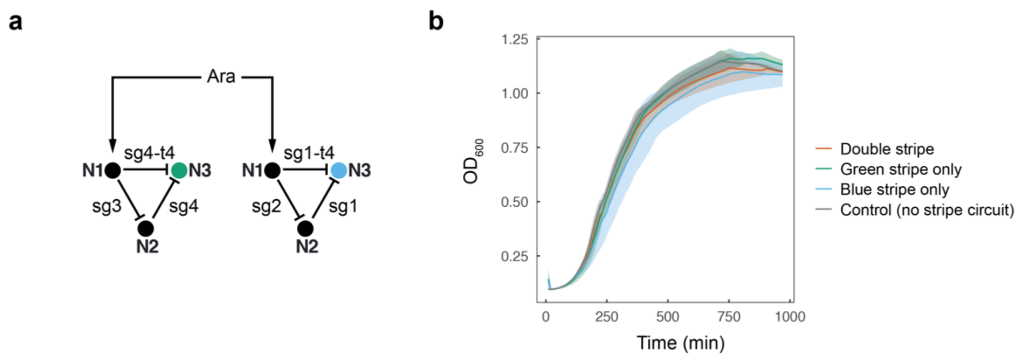


**Supplementary Fig. 3.** Increase of CRISPRi circuit complexity does not impact cell growth. (**a**) Design of the double stripe-forming two-IFFL circuit. (**b**) Growth curves of bacteria carrying the double stripe circuit depicted in (a) compared to cells carrying either the blue stripe or the green stripe circuits only, or cells lacking both stripe networks. Lines and shades correspond to the mean and s.d. of 3 biological replicates, respectively.

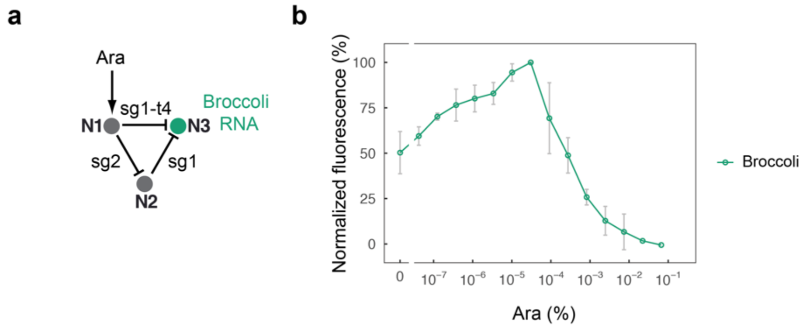


**Supplementary Fig. 4.** A RNA-reported and RNA-controlled IFFL. (**a**) Design of the synthetic circuit. (**b**) Stripe behavior of the CRISPRi IFFL as reported by a Broccoli RNA aptamer in the presence of DFHBI-1T. Mean and s.d. of three biological replicates.

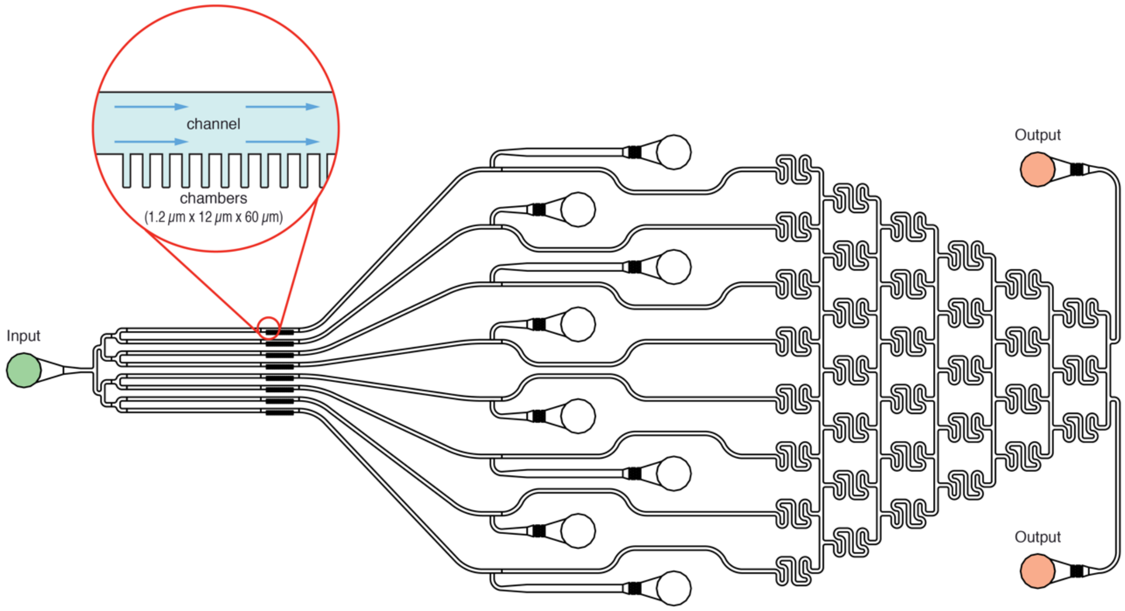


**Supplementary Fig. 5.** Design of the microfluidic device. Bacteria were grown in chambers (1.2 µm x 12 µm x 60 µm, h x w x l) harboring approximately 110 cells each. The downstream, tree-like design can be used to generate a concentration gradient. This feature was not used in this study.

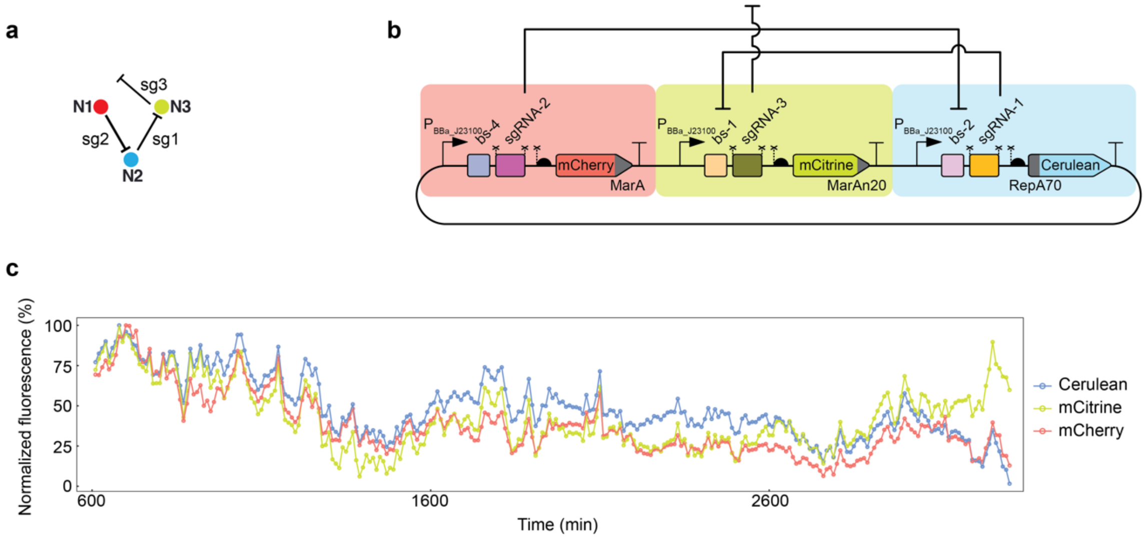


**Supplementary Fig. 6.** An "open-ring" control for the CRISPRlator lacking any oscillatory behavior. (**a**) "Open-ring" topology of the control circuit. (**b**) Molecular implementation. The sgRNA-4 present in the CRISPRlator was replaced with sgRNA-3, which has no cognate binding site in the circuit and is thus unable to "close" the repression ring. Symbols as in Fig. 1. (**c**) Quantification of the population-level fluorescence over time.

| Abbreviation | Explanation |
|---|---|
| G1 | Gene 1 |
| G2 | Gene 2 |
| G3 | Gene 3 |
| G4 | Gene 4 |
| sg1 | single guide RNA 1 |
| sg2 | single guide RNA 2 |
| Cas | dCas9 complex |
| Cassg1 | Cas with sg1 |
| Cassg2 | Cas with sg2 |
| G1P2 | Cassg2 specifically bound to G1 |
| G2P1 | Cassg1 specifically bound to G2 |
| G3P1 | Cassg1 specifically bound to G3 |
| G4P2 | Cassg2 specifically bound to G4 |
| G1U1 | Cassg1 unspecifically bound to G1 |
| G1U2 | Cassg2 unspecifically bound to G1 |
| G2U1 | Cassg1 unspecifically bound to G2 |
| G2U2 | Cassg2 unspecifically bound to G2 |
| G1P2U1 | Cassg2 specifically and Cassg1 unspecifically bound to G1 |
| G1P2U2 | Cassg2 specifically and Cassg2 unspecifically bound to G1 |
| G2P1U1 | Cassg1 specifically and Cassg1 unspecifically bound to G2 |
| G2P1U2 | Cassg1 specifically and Cassg2 unspecifically bound to G2 |

**Supplementary Table 2.** Abbreviations and functionality of the species used in models of the CRISPRi toggle switch (TS). Note that sgRNA-1 (abbreviated sg1) and sgRNA-2 (sg2) in the model represent two “generic” sgRNAs, and not necessarily the specific sequences labelled as “sgRNA-1” and “sgRNA-2” in the experimental parts of this work.

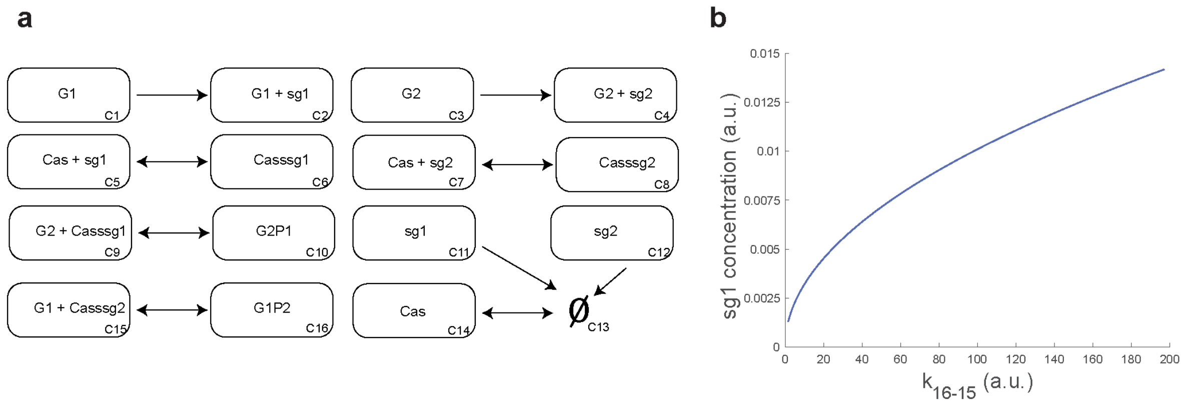


**Supplementary Fig. 7.** Simplest model for the TS (case 1). (**a**) Model structure assuming that Cassg1/2 complexes bind to a single, specific site on their target genes. Boxes denote complexes in chemical reaction network theory, that is, subsets of reactants (Supplementary Table 2) that are jointly educts or products of a reaction (complex identifiers in lower right corner). Single-headed arrows: irreversible reactions; double-headed arrows: reversible reactions; Ø: source or sink for components. (**b**) Bifurcation diagram of the steady state sg1 concentration upon varying the constant $k_{16\text{-}15}$, which corresponds to the reaction of complexes C16 and C15 (unbinding of the specifically bound Cassg2 to G1). The analysis indicates that the system cannot show bistability.

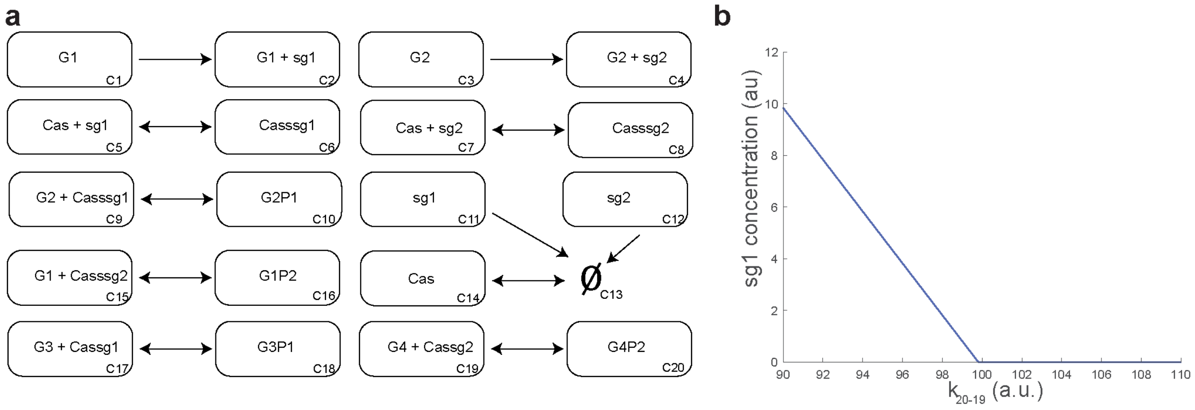


**Supplementary Fig. 8.** Extended model for the TS (case 2). (**a**) Model structure, assuming that the Cassg1/2 complexes can bind to other sequences in the genome specifically and not only to G1 and G2. Notation as in Supplementary Fig. 7. (**b**) Bifurcation diagram of the steady state sg1 concentration with varying the constant $k_{20\text{-}19}$, which corresponds to the reaction of complexes C20 and C19 (unbinding of the specifically bound Cassg2 to G4).

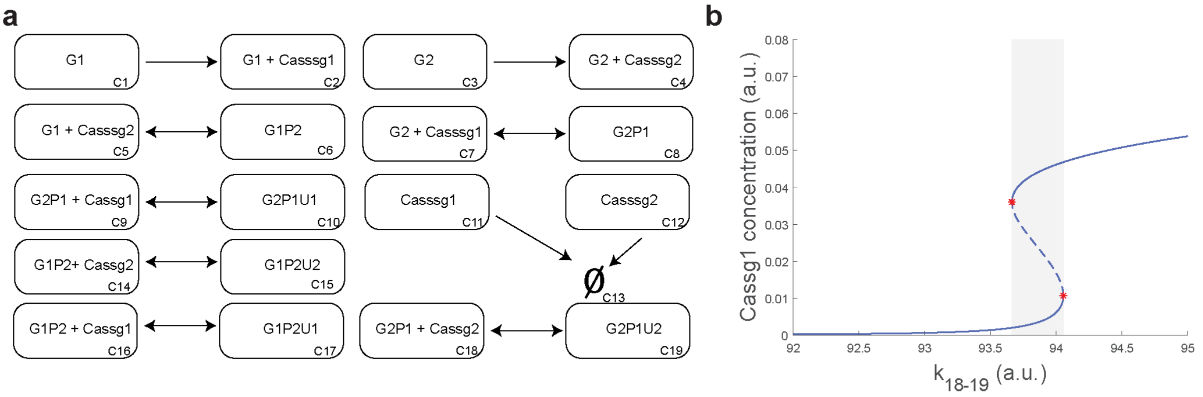


**Supplementary Fig. 9.** Model for the TS with unspecific binding (case 3). (**a**) Model structure as in Supplementary Fig. 7. The model assumes that we can neglect the formation of the Cassg1/2 complexes and that the Cassg1/2 complexes can bind unspecifically via PAM sequences to G1 and G2. (**b**) Bifurcation diagram of the steady state Cassg1 concentration with varying the constant $k_{18\text{-}19}$, which corresponds to the reaction of complexes C18 and C19 (unspecific binding of Cassg2 to G2P1). Stable and unstable steady states are represented by solid and dashed lines, respectively. Red stars indicate limit points and bistability regions are enclosed in the grey area.

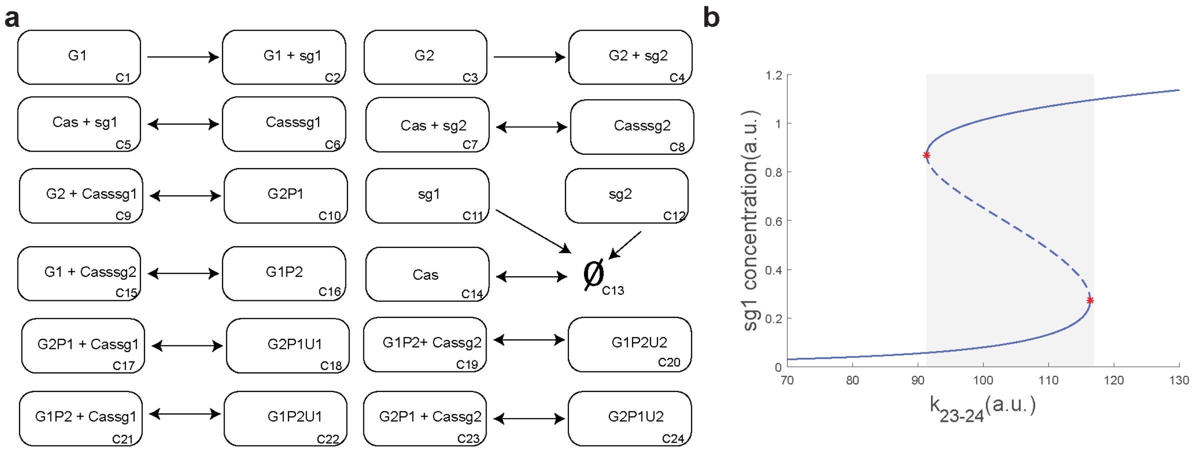


**Supplementary Fig. 10.** Model for the TS with unspecific binding (case 4). (**a**) Model structure, extension of case 3 (Supplementary Fig. 9) by Cassg1/2 complex formation. (**b**) Bifurcation diagram of the steady state sg1 concentration with varying the constant $k_{23\text{-}24}$, which corresponds to the reaction of complexes C23 and C24 (unspecific binding of Cassg2 to G2P1). Symbols are as in Supplementary Fig. 9.

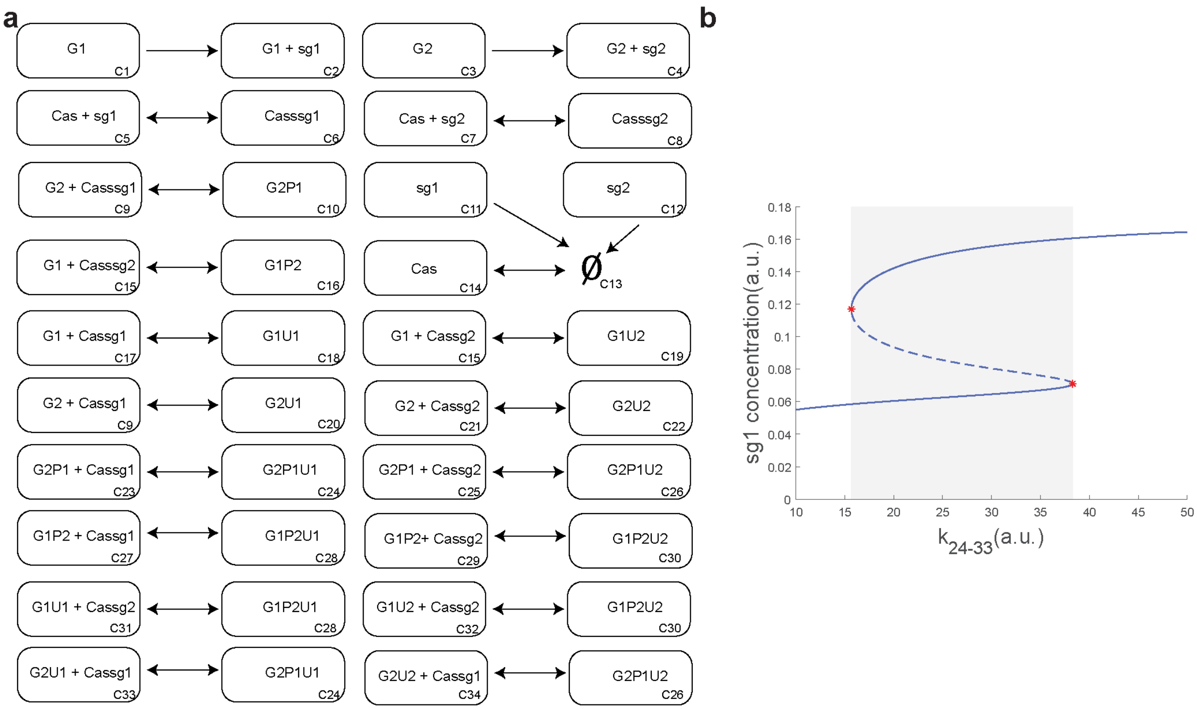


**Supplementary Fig. 11.** Model for the TS with unspecific binding (case 5). (**a**) Model structure, which extends case 4 (Supplementary Fig. 10) to include the unspecific binding of Cassg1 and Cassg2 to G2 and G1, respectively. (**b**) Bifurcation diagram of the steady state concentration of sg1 with varying constant $k_{24\text{-}33}$, which corresponds to the reaction of complexes C24 and C33 (unbinding of the unspecific complex Cassg1 from G2P1U1). Symbols are as in Supplementary Fig. 9.

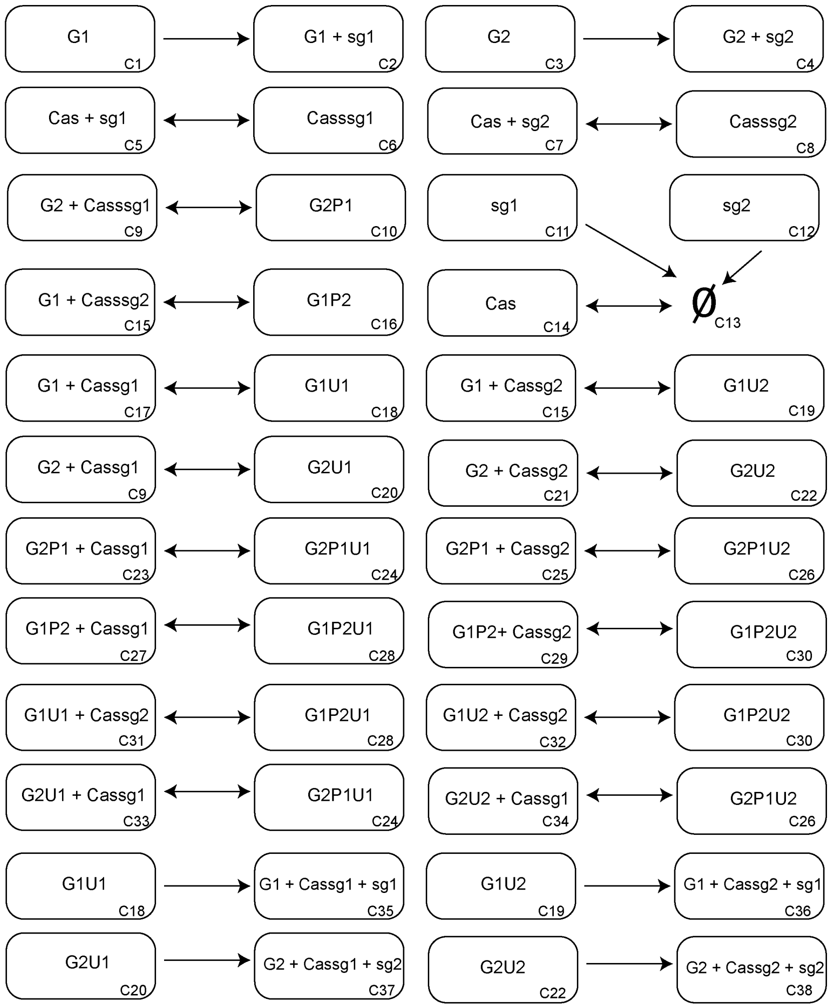


**Supplementary Fig. 12.** Complete model for the TS. Compared to the other cases, this model assumes that unspecific binding of Cassg1/2 complexes does not affect gene expression. The bifurcation diagram of this model with experimentally constrained parameters (Supplementary Table 3) is shown as Fig. 3b in the main text.

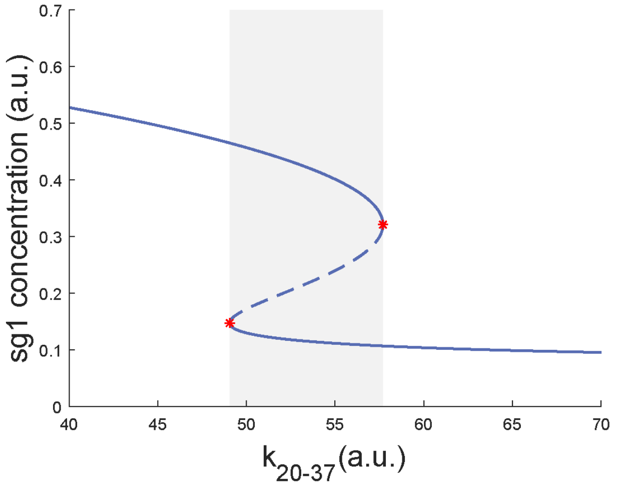


**Supplementary Fig. 13.** Bifurcation diagram of the complete model without experimentally-derived constraints on parameters. The varying parameter corresponds to the reaction of complexes C20 and C37 (Supplementary Fig. 12) and is similar to the one represented in Fig. 3b (main text). Symbols are as in Supplementary Fig. 9.

| Parameters | Process | Lower Bound | Upper Bound | pBest |
|---|---|---|---|---|
| k_1_2 | Production | 1 | 100 | 50.72434 |
| k_3_4 | Production | 1 | 100 | 42.33746 |
| k_13_14 | Production | 1 | 100 | 1.031285 |
| k_18_35 | Production | 0.1 | 100 | 65.26698 |
| k_19_36 | Production | 0.1 | 100 | 75.94251 |
| k_20_37 | Production | 0.1 | 100 | 68.53327 |
| k_22_38 | Production | 0.1 | 100 | 63.44098 |
| k_5_6 | Association | 1 | 100 | 1.188162 |
| k_7_8 | Association | 1 | 100 | 1.071073 |
| k_6_5 | Dissociation | 0.01 | 10 | 9.601046 |
| k_8_7 | Dissociation | 0.01 | 10 | 9.432942 |
| k_11_13 | Degradation | 0.001 | 1 | 0.986721 |
| k_12_13 | Degradation | 0.001 | 1 | 0.964785 |
| k_14_13 | Degradation | 0.001 | 1 | 0.988306 |
| k_9_10* | Specific binding | 1 | 100 | 4.612951 |
| k_15_16* | Specific binding | 1 | 100 | 33.02472 |
| k_31_28* | Specific binding | 1 | 100 | 42.83338 |
| k_32_30* | Specific binding | 1 | 100 | 76.30581 |
| k_33_24* | Specific binding | 1 | 100 | 51.87789 |
| k_34_26* | Specific binding | 1 | 100 | 65.45974 |
| k_10_9* | Specific unbinding | 0.001 | 0.1 | 0.093696 |
| k_16_15* | Specific unbinding | 0.001 | 0.1 | 0.096394 |
| k_28_31* | Specific unbinding | 0.001 | 0.1 | 0.089324 |
| k_30_32* | Specific unbinding | 0.001 | 0.1 | 0.094013 |
| k_24_33* | Specific unbinding | 0.001 | 0.1 | 0.085184 |
| k_26_34* | Specific unbinding | 0.001 | 0.1 | 0.095339 |
| k_17_18* | Unspecific binding | 1 | 100 | 76.13001 |
| k_15_19* | Unspecific binding | 1 | 100 | 94.13042 |
| k_9_20* | Unspecific binding | 1 | 100 | 85.43037 |
| k_21_22* | Unspecific binding | 1 | 100 | 94.57936 |
| k_23_24* | Unspecific binding | 1 | 100 | 53.13512 |
| k_25_26* | Unspecific binding | 1 | 100 | 86.54855 |
| k_27_28* | Unspecific binding | 1 | 100 | 89.51009 |
| k_29_30* | Unspecific binding | 1 | 100 | 65.04785 |
| k_18_17* | Unspecific unbinding | 1 | 100 | 11.96749 |
| k_19_15* | Unspecific unbinding | 1 | 100 | 3.707992 |
| k_20_9* | Unspecific unbinding | 1 | 100 | 12.95706 |
| k_22_21* | Unspecific unbinding | 1 | 100 | 5.169212 |
| k_24_23* | Unspecific unbinding | 1 | 100 | 1.177815 |
| k_26_25* | Unspecific unbinding | 1 | 100 | 1.521193 |
| k_28_27* | Unspecific unbinding | 1 | 100 | 1.472313 |
| k_30_29* | Unspecific unbinding | 1 | 100 | 1.446816 |

**Supplementary Table 3.** Parameters for the complete model with experimental constraints. Parameter names represent the reaction between the complexes in which they participate; 'k_i_j' is the rate constant for the irreversible reaction from complex $C_i$ to $C_j$. During parameter space exploration for identifying a limit point, parameters denoted with * were constrained in a range defined by experimental data from Martens *et al.* 2019, Nat. Commun. 10, 3552. Specifically, we allowed for +/- two orders of magnitude for unspecific binding and +/- one order of magnitude for specific binding. pBest is the optimal decision vector.
