## Supplementary material for "Multistable and dynamic CRISPRi-based synthetic circuits": Code for model analysis: case2.m

```
function [ESP, COM, KNS] = case2
% Case 2 described in supplementary: assuming specific binding to G1, G2,
% G3, G3

ESP={'G1','sg1','G2','sg2','Cas','Cassg1','Cassg2','G2I','G1I','G3','G3I','G4','G4I'};

COM={'G1','G1 + sg1','G2','G2 + sg2','Cas + sg1','Cassg1','Cas + sg2','Cassg2', ...
    'G2 + Cassg1','G2I','G1 + Cassg2','G1I','sg1','sg2','0','Cas',...
    'Cassg1 + G3','G3I','Cassg2 + G4','G4I'}; 


KNS={'k_1_2','k_3_4','k_5_6','k_6_5','k_7_8','k_8_7','k_9_10','k_10_9','k_11_12','k_12_11', ...
    'k_13_15','k_14_15','k_15_16','k_16_15','k_17_18','k_18_17','k_19_20','k_20_19'};

end
```
