## Supplementary material for "Multistable and dynamic CRISPRi-based synthetic circuits": Code for model analysis: case3.m

```
function [ESP, COM, KNS] = case3
% Case 3 from supplementary: removing dCas formation,simple unspecific
% binding

ESP={'G1','P1','G2','G1P2','G2P1','G2P1P1','P2','G1P2P2','G1P2P1','G2P1P2'};

COM={'G1','G1+P1','G2','G2+P2','G1+P2','G1P2','G2+P1','G2P1','G2P1+P1','G2P1P1','P1','P2','0',...
    'G1P2 + P2', 'G1P2P2', 'G1P2 + P1', 'G1P2P1', 'G2P1 + P2', 'G2P1P2'}; 

% reversible Cas9 binding
KNS={'k_1_2' 'k_3_4' 'k_5_6' 'k_6_5' 'k_7_8' 'k_8_7'  'k_9_10' 'k_10_9' ...
    'k_11_13' 'k_12_13'  'k_14_15' 'k_15_14'  'k_16_17' 'k_17_16' 'k_18_19' 'k_19_18'};

end
```
