## Supplementary material for "Multistable and dynamic CRISPRi-based synthetic circuits": Code for model analysis: case4.m

```
function [ESP, COM, KNS] = case4
% Case 4 from supplementary- adding dCas formation and simple case of unspecific binding

ESP={'G1','P1','G2','G1P2','G2P1','G2P1P1','P2','G1P2P2','G1P2P1','G2P1P2','S1','S2','C0'};

COM={'G1','G1 + S1','G2','G2 + S2',...
    'C0 + S1','P1','C0 + S2','P2',...
    'G2 + P1','G2P1',...
	'S1','S2','0','C0',...
    'G1 + P2','G1P2',...
    'G2P1+P1','G2P1P1','G1P2 + P2', 'G1P2P2','G1P2 + P1', 'G1P2P1', 'G2P1 + P2', 'G2P1P2' };
 

KNS={'k_1_2' 'k_3_4' 'k_5_6' 'k_6_5' 'k_7_8' 'k_8_7'  'k_9_10' 'k_10_9' ...
    'k_11_13' 'k_12_13'  'k_14_13' 'k_13_14'  'k_15_16' 'k_16_15' 'k_17_18' 'k_18_17'...
    'k_19_20' 'k_20_19'  'k_21_22' 'k_22_21' 'k_23_24' 'k_24_23'};
end
```
