## Supplementary material for "Multistable and dynamic CRISPRi-based synthetic circuits": Code for model analysis: case6.m

```
function [ESP, COM, KNS] = case6
%Case 6 from supplementary: unspecific binding not interfering with
%transcription


ESP={'G1','G2','S1','S2','C0S1','C0S2','C0','G2P1','G1P2','G2U1','G2U2','G1U1','G1U2','G1P2U1','G1P2U2','G2P1U1','G2P1U2'};

COM={'G1','G1 + S1',...
    'G2','G2 + S2',...
    'C0 + S1','C0S1',...
    'C0 + S2','C0S2',...
    'G2 + C0S1','G2P1',...
	'S1','S2','0','C0',...
    'G1 + C0S2','G1P2',...
    'G1 + C0S1','G1U1',...
    'G1U2',...
    'G2U1',...
    'G2 + C0S2','G2U2',...
    'G2P1 + C0S1','G2P1U1',...
    'G2P1 + C0S2','G2P1U2',...
    'G1P2 + C0S1','G1P2U1',...
    'G1P2 + C0S2','G1P2U2',...
    'G1U1 + C0S2',...
    'G1U2 + C0S2',...
    'G2U1 + C0S1',...
    'G2U2 + C0S1',...
    'G1 + C0S1 + S1',...
    'G1 + C0S2 + S1',...
    'G2 + C0S1 + S2',...
    'G2 + C0S2 + S2'};
    

KNS={'k_1_2' ,'k_3_4' ,'k_5_6', 'k_6_5', 'k_7_8', 'k_8_7',  'k_9_10', 'k_10_9', ...
    'k_11_13' ,'k_12_13',  'k_13_14', 'k_14_13',  'k_15_16', 'k_16_15',...
    'k_17_18', 'k_18_17' ,'k_15_19', 'k_19_15' , 'k_9_20' ,'k_20_9', 'k_21_22' ,'k_22_21',...
    'k_23_24','k_24_23','k_25_26','k_26_25','k_27_28','k_28_27','k_29_30','k_30_29',...
    'k_31_28','k_28_31','k_32_30','k_30_32','k_33_24','k_24_33','k_34_26','k_26_34',...
    'k_18_35','k_19_36','k_20_37','k_22_38'};
end
```
