## Supplementary material for "Multistable and dynamic CRISPRi-based synthetic circuits": Code for model analysis: README.txt

Intro
=====

For identifying the nonlinear behaviour of the toggle switch using CRISPRi constructs, we employed BioSwitch [1]. BioSwitch is a Matlab software for detecting multistable biochemical systems.
The code provided includes all the model structures we evaluated (case1-case6) and a script with the necessary commands to use with BioSwitch.


Installation
============

Download and install BioSwitch [1], MEIGO optimization toolbox [2] and MatCont [3]. 


Quickstart
==========

First take a look into the `run_case1_example` script to see the first model that was tested to generate bistability.

1. To be able to run `run_case1_example` it should be in the BioSwitch path (see BioSwitch documentation).

2. Each command in the script generates the nesseccary files to evaluate multistability.

3. To test all the mentioned models replace with the corresponding name function.


Authors
=======

    Santos-Moreno, Javier <>>
    Tasiudi, Eve <>
    Stelling, Joerg <>>
    Schaerli, Yolanda <>>


References
==========

[1]: https://github.com/ireneotero/BioSwitch
[2]: http://gingproc.iim.csic.es/meigom.html
[3]: https://sourceforge.net/projects/matcont/files/Oldcl_matcont/cl_matcont5p4/
