## Supplementary material for "Multistable and dynamic CRISPRi-based synthetic circuits": Code for model analysis: run_case1_example.m

```
% script to run with Bioswitch for the different cases
% case 1: simple case
% case 2: assuming specific binding to G1, G2, G3 and G4
% case 3: removing dCas formation, simple unspecific binding
% case 4: adding dCas formation, simple unspecific binding
% case 5: full model with unspecific binding (unspecific binding interferes
% with transcription)
% case 6: complete model (unspecific binding does not interfere with
% transcription)

% To test each model structure, replace with the corresponding case to be
% tested.

% Evaluate Network
BioSWITCH_Evaluate('case1')

% Generate files for limit point search (here system with mass conservation
% was used)
BioSWITCH_mkfiles_lpsearch_MassCon('case1');

% Search for limit point
BioSWITCH_Lpsearch('case1_input_lpsearch_MassCon_default');

% Create files for equlibrium continuation
% The vector must be changed depending on the conservation relations in each case
BioSWITCH_mkfiles_ncontin_MassCon('case1', [1,3]) 

% Perform continuation
BioSWITCH_Ncontin('case1_input_cont_MassCon_default');
```
